# Multimodal AI for Single cfDNA Profiling and Cancer Screening

**DOI:** 10.64898/2025.12.29.696856

**Authors:** Bo Wang, Liyang Song, Hefei Li, Nan Lin, Ying Xin, Xiaowen He, Wenxin Liu, Li Liu, Jian Cui, Xuesong Li, Ying Mei, Qiuting You, Haodong Zhu, Guoqiang Zhao, Guo Chen, Jing Liu, Baoliang Zhu, Xueguang Sun, Xiaohui Wu, Zhidong Gao, Yingjiang Ye

## Abstract

Cell-free DNA (cfDNA) serves as a non-invasive biomarker for cancer detection, but conventional methods face challenges due to the ultra-low abundance of tumor-derived cfDNA (ctDNA) among normal cfDNA. Though nucleosome-bound cfDNA harbors rich epigenomic features that could enable ctDNA identification by single-molecule multi-omics cross-validation, this remains unexplored due to methodological limits. Here, we developed a cfDNA sequencing approach integrating methylation, fragmentomics, and histone modifications at the single-molecule level; together with gene semantics and epigenomic annotations, these modalities were vectorized and fused to represent each cfDNA molecule. We trained a Transformer-based model (cfAI) to profile and evaluate ctDNA likelihood at molecule, gene, and sample levels. cfAI achieved ∼10-fold enrichment of cancer-derived signals over noise and reached 72.6% sensitivity at 93.1% specificity for multi-cancer detection. Our study establishes an innovative framework that overcomes the inherent signal-to-noise limitations of conventional assays and reveals biological features at molecular resolution for cancer detection.

## Introduction

Cell-free DNA (cfDNA) originates from dying cells (apoptosis, necrosis, etc.) and carries molecular information reflective of the physiological state of its tissues of origin^1–3^. Once released into the circulation, cfDNA can be noninvasively extracted and used to monitor changes in tumors, the fetus, transplanted organs, and other sources^4–6^. Tumor-derived cfDNA (circulating tumor DNA, ctDNA) comprises a heterogeneous set of abnormalities spanning genomic mutations, epigenomic alterations, fragmentomic features, and histone modifications—i.e., multi-omics signals^7–11^. A variety of cfDNA-sequencing–based noninvasive diagnostic and screening approaches are now well established and have achieved both technical and commercial success^12–14^. A major challenge in early cancer screening using ctDNA is its extremely low abundance, often less than 0.1%^15,16^, which is easily overshadowed by background cfDNA from normal cells^17,18^. As a result, those ctDNA can only be estimated based on subtle fluctuations in epigenomic values within the overall cfDNA mixture. This mixture-based strategy hampers every step, including both biomarker screening and diagnostic assay measurement^19^. Consequently, existing examples typically rely on markers that require clean background signals or linkage across CpGs to form detectable haplotype^20,21^, with few markers (like SEPT9)^22^ able to work in such a noisy environment. Other methods attempt to overcome this by smoothing signals over larger genomic windows (megabase and arm levels)^4,23,24^, but the actual variation often occurs in much smaller regions, such as specific nucleosome positions^9,25^. Therefore, mixture-based methods struggle to detect signals with sufficient accuracy at affordable sequencing depths (ten-thousand–fold noise vs. thousands-fold depth), which remains the primary bottleneck in advancing next-generation early cancer screening^26–28^.

Notably, a low ctDNA fraction does not mean it is indistinguishable: cfDNA is released as nucleosomes, while multiple signals co-localize on the same physical unit. Studies have shown that these multi-omic features act synergistically^25^ and are closely linked to gene expression, transcription-factor binding, 3D genome organization, and chromatin state^1,9^. The ultimate goal is to characterize multiple omic modalities simultaneously at single-cfDNA-molecule resolution and exploit their cross-relationships to substantially increase signal-to-noise. Modeling each omic separately and then fusing results sacrifices molecule-level granularity and loses cross-feature information.

Unfortunately, most cfDNA pipelines require preformatted `<feature_n>: <value>` pairs^29,30^; forcing mixture-based aggregation and preventing retention of per-molecule feature sets and their cross-feature / cross-molecule interactions.

The recent surge in Transformer-based AI models has proven powerful for modeling complex, high-dimensional data^31,32^. Recent advances in single-cell and multi-omic large models offer useful architectural templates and prior knowledge^33,34^. However, those models treat words, cells, or genes as tokens — cfDNA requires treating each molecule as a token and fusing many heterogeneous features. Therefore, accurately modeling cfDNA requires embedding not only the raw omic feature values but also the corresponding genomic coordinates and gene-level semantic context.

In this study, we developed a new experimental method to capture multiple orthogonal omic signals—including DNA methylation, fragmentomic features, and histone modifications—on the same cfDNA molecule. Building on this, we pioneer multimodal integration at single-cfDNA-molecule resolution to characterize synergistic multi-omic changes associated with cancer, enabling ctDNA-level inference, gene-level prediction, and sample-level diagnosis. This proof-of-concept demonstrates the feasibility of the approach and advances the frontier of liquid biopsy and early cancer screening.

## Results

### Study Design of cfAI

We developed a multi-omic sequencing protocol to capture multiple orthogonal signals on the same cfDNA molecule (Fig. 1). Plasma from peripheral blood was subjected to antibody-conjugated magnetic bead pull-down to enrich nucleosome complexes. By targeting H3K4me3 and focusing capture near transcription start sites (TSS), we enriched nucleosomes associated with high transcriptional activity, thereby amplifying differential signals. The enriched cfDNA was eluted and purified, followed by nick-repaired, dual-strand library construction and enzymatic methylation conversion. These steps preserve fragment end integrity in highly nicked cfDNA and protect 5mC near 3’ ends from polymerase erasing (see Methods and Fig. S1).

**Figure 1.**
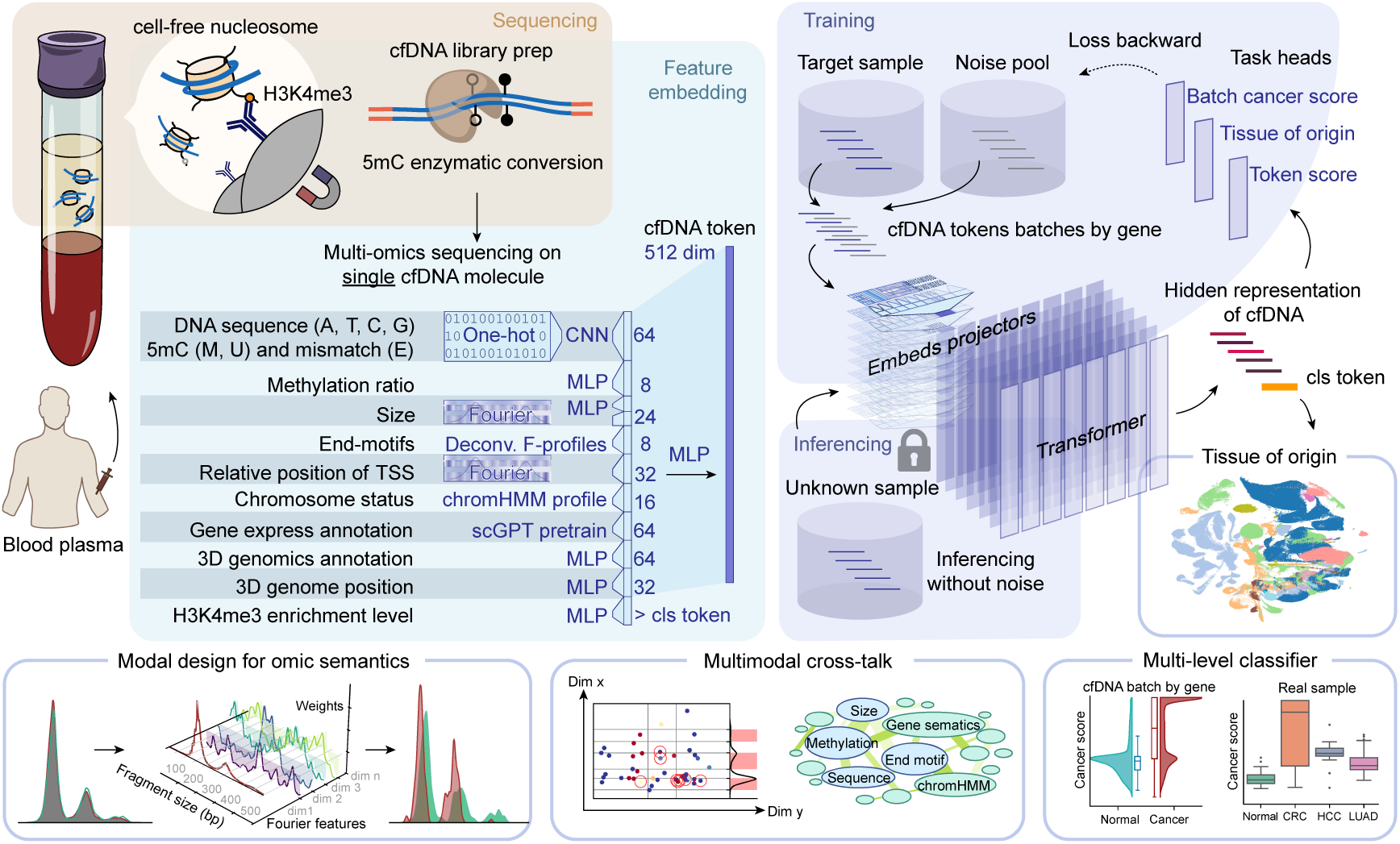
Schematic of study design. cf-nucleosomes in plasma were captured with H3K4me3-affinity antibody-coated magnetic beads and subjected to an enzymatic methylation-conversion sequencing workflow that preserves multi-omic co-information. For each cfDNA fragment (read), multiple omic features were extracted by bioinformatic processing and mapped into a tailored vector-embedding representation. The multimodal representations were merged into a single cfDNA token. During model training, the embedding layer, Transformer layers, and multi-task heads are trained end-to-end. The learnable embedding layer can attend to multiple aspects of each omic modality and produce semantically rich encodings; the multimodal fusion layer and Transformer layers enable crosstalk among modalities. Model outputs include per-cfDNA scores, cfDNA gene-batch–level diagnoses, and information-rich latent states.

For each cfDNA fragment we extracted multi-modal features including DNA sequence, CpG methylation sites and methylation ratios, mismatches, fragment length, and end motifs. Fragments were further annotated with high-resolution regulatory context such as 3D genome features (e.g., A/B compartments), chromatin segmentation (e.g., ChromHMM states^35^), and gene semantics from pretrained models^34^.

Each feature type was projected into embedding vectors using modality-specific mappers designed based on underlying biology and data structure—for example, learnable Fourier feature encodings, convolutional networks (CNN) for sequence patterns, and emission-matrix followed by UMAP for certain regulatory signals. The per-feature embeddings were concatenated and passed through multi-layer perceptrons (MLPs) to produce a single latent vector for each cfDNA fragment, which we treat as a token.

We trained a Transformer-based model that takes cfDNA tokens as input and captures both intra-molecular features and inter-molecular relationships via attention. During pretraining, tokens from target samples were mixed with noise tokens sampled from an equal-depth pool of healthy cfDNA. Tokens were grouped into cfDNA batches by their corresponding gene. Sequencing depth and H3K4me3 enrichment was packed into a <cls> token. The model is trained to predict tissue-of-origin at the cfDNA-level and at the cfDNA-batch level. At inference, inputs are provided without noise mixing, and the cancer score of the sample is computed as the mean of its cfDNA-batch cancer scores.

We recruited 333 participants (including 10 cancer types) across 11 clinical sites; 304 qualified samples were randomly split into training (n = 180) and testing (n = 124) sets. Cutoffs were determined on the training set and evaluated on the locked test set (see Methods and Fig. S2).

By combining a sequencing protocol that jointly captures methylation, fragmentomic, and histone signals with an end-to-end Transformer trained on per-molecule embeddings, we present a high-resolution multimodal fusion framework at single-cfDNA-molecule resolution. This architecture enables integrated use of heterogeneous cfDNA signals for ctDNA inference, gene-level prediction, and sample-level discrimination.

### Single-molecule multi-omic synergy enables effective ctDNA enrichment

We observed clear multi-omic co-occurrence localized on individual cfDNA molecules. When projecting each cfDNA as a point in a multi-feature space, tumor-derived fragments form distinct patterns and occupy specific regions of that space. Taking IRS1, a well-characterized cancer gene, as an example and jointly examining nucleosome position, fragment length, and CpG methylation (Fig. 2A; x-axis, y-axis, color bar), tumor cfDNA shows an excess of low-methylation, short-fragment molecules that cluster at particular nucleosome positions—notably within promoter regions (Fig. 2A, lower panel). This pattern is concordant with IRS1’s tumor-specific overexpression and its association with poor prognosis^36^. Thus, because of pronounced multi-omic synergy, ctDNA can be readily identified by integrating cross-dimensional signals. By contrast, unimodal mixture statistics miss this feature: fragment-only comparison of the short-fragment fraction shows no difference between cancer and normal (p = 0.3, Mann–Whitney U). Meanwhile, selecting fragments by methylation and nucleosome position dramatically amplifies the short-fragment signal (p = 0.007) (Fig. 2B), confirming that single-omic analyses waste cross-modal information and miss valuable markers. Scoring fragments with our single-molecule multi-omic cfAI model highlights these specific loci (Fig. 2A) and further enriches ctDNA features—raising fragment significance to p = 0.00026 (Fig. 2B) and markedly amplifying low-methylation signal within the differential-methylation region (Fig. 2C). These results indicate that the cfAI model correctly identifies and enriches ctDNA.

**Figure 2.**
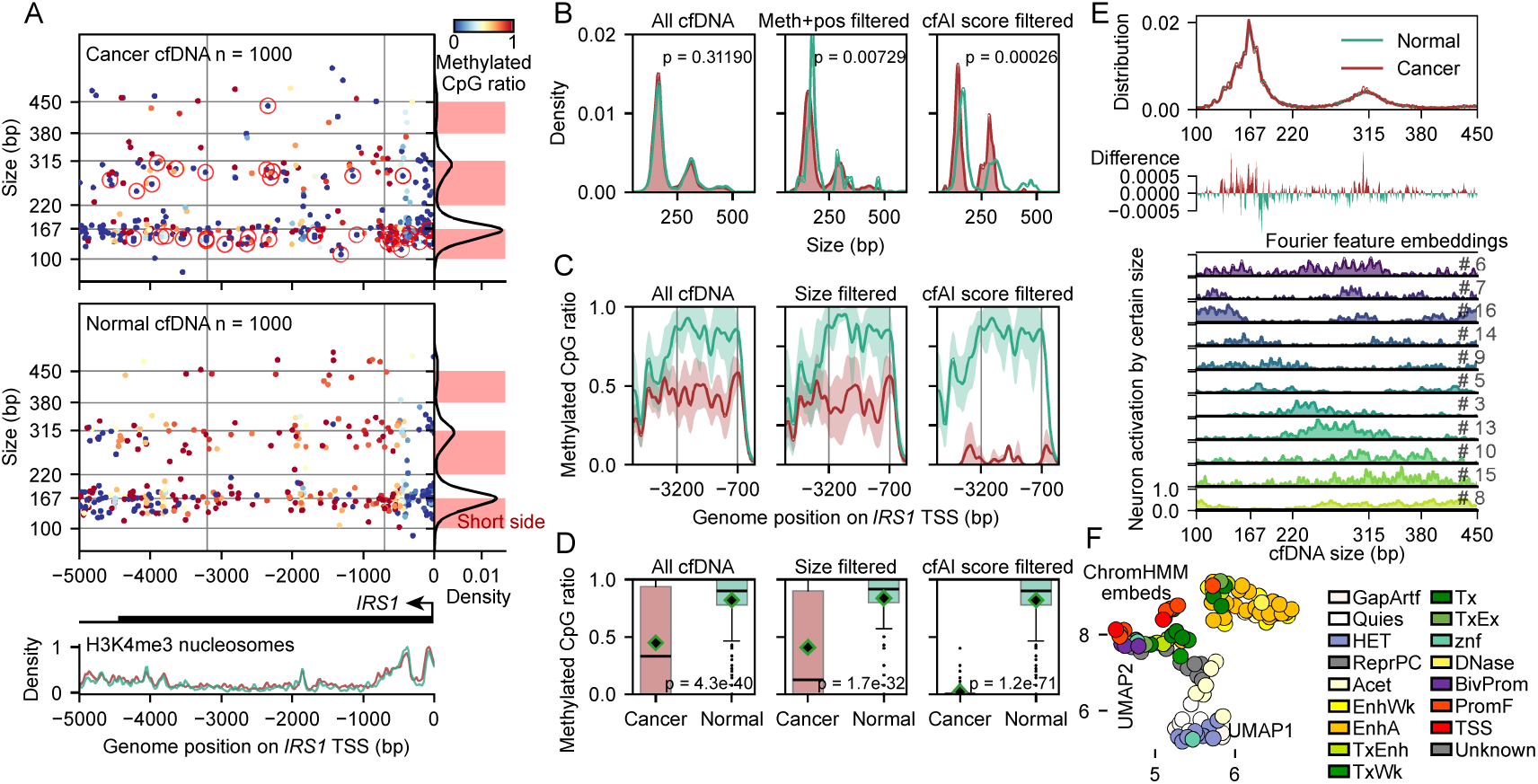
Single-molecule multi-omic synergy enables effective ctDNA enrichment. (A) ctDNA is identified intuitively by comparing multi-omics data of cancer and normal cfDNA in the IRS1 promoter region. Each point represents a cfDNA molecule; the x-axis represents the fragment’s relative position to the IRS1 TSS, the y-axis shows fragment length, and color encodes CpG methylation level. Red circles denote ctDNA identified by cfAI. Marginal density plots (bottom and right) indicate nucleosome occupancy and fragment-length distributions; the peak side enriched for shorter fragments is highlighted in red. (B–D) Distributions of fragment length and methylation across all cfDNA at this locus (left), their amplification after multi-omic filtering (middle), and after cfAI selection (right). Fragmentomic statistics used the short-side fraction and were tested with the Mann–Whitney U test (B). Methylation plots display the DMR region with cancer and normal methylation curves and a 95% confidence CI (shaded); the DMR methylation distribution was quantified in (C). (E) Subtle fragmentomic differences in cancer and the attention of different neurons to fragment-length features; an example neuron detects a 10.7-bp periodicity. (F) Two-dimensional UMAP of chromHMM embedding vectors, illustrating unsupervised clustering that reflects the model’s learned genomic context.

From the fragmentomic example, we note that both long and short fragments can be tumor-derived; the relevant signal is whether a fragment lies on the short- or long-side of mono-, di- or tri-nucleosome distribution peaks (Fig. 2A, red region), rather than an absolute scalar such as “fragment length = 170 bp.” To capture such complex relationships, we designed modality-specific mappings that combine domain priors, custom activation functions, dimensional scaling, and end-to-end learning. For fragment length, we decomposed semantics with learnable Fourier features (wavelength and phase), enabling different neurons to attend to different periodicities—one neuron (Neuron #6) recovers a 10.7-bp periodicity on the short-fragment side (Fig. 2E), consistent with nuclease digestion preferences reflecting DNA wrapping topology and a known ctDNA signal. We also incorporated high-quality prior annotations—gene semantics from single-cell large models and a trained chromHMM functional annotation—to produce genomically contextualized embeddings with biological interpretability (Fig. 2F), instead of relying solely on raw nucleotide strings.

### Context-aware modeling across multiple cfDNA fragments enhances denoising and discrimination

We designed cfAI to process, as a single input batch, all upstream and downstream cfDNA molecules from the same sample and gene, enabling joint inference across that context. The model outputs both a per-fragment cancer propensity and a denoised batch-level cancer score. Consistent with a very low ctDNA fraction, most individual cfDNA fragments receive unremarkable cancer scores and—even when tumor-derived—have average scores like normal fragments. However, by aggregating contextual information across the whole gene-level batch, the model extracts signal from these low means and produces elevated batch-level scores (Fig. 3A, lower right); this improvement is also confirmed on the held-out test set.

**Figure 3.**
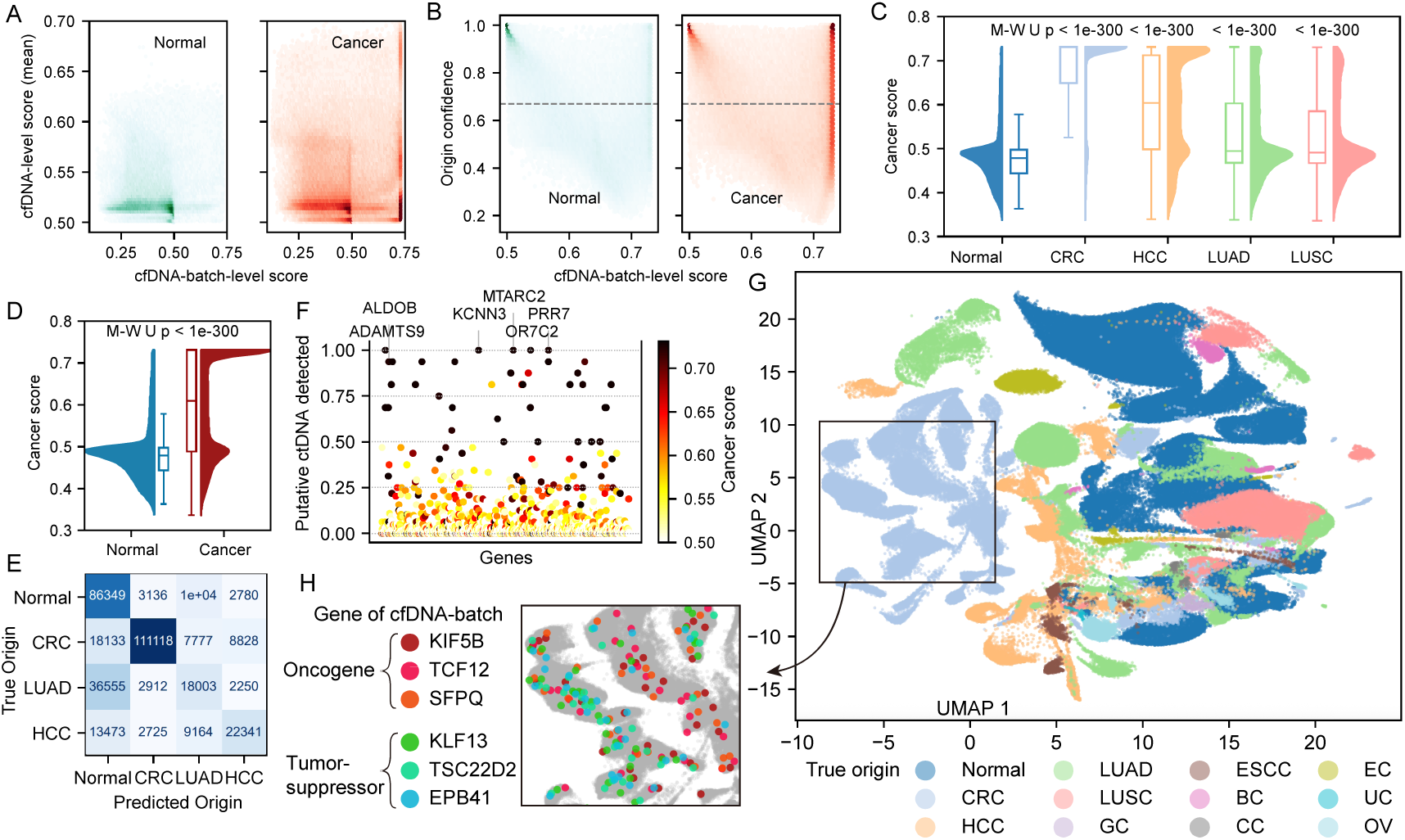
Context-aware aggregation of multiple cfDNA fragments improves denoising and discrimination by biology representations. (A) Relationship between the mean per-fragment cancer score (vertical axis) and the aggregated batch-level score (horizontal axis) for cfDNA batches. Many cancer batches have low mean per-fragment scores but show a marked increase in aggregated score (lower right), indicating effective denoising by the model. (B) Relationship between aggregated score (horizontal axis) and origin-prediction confidence (vertical axis); the dashed line denotes the confidence cutoff used for downstream gating. (C–D) Comparison of gated cancer scores between normal and individual cancer types (C), and normal versus all cancer types (D) using the Mann-Whitney U test (P< 1×10−^308^); (E) Confusion matrix for cfDNA-batch origin prediction. (F) Genes that were stably detected as putative ctDNA across multiple batches; color encodes the gene-wise average cancer score. (G) Two-dimensional UMAP of CLS hidden-state embeddings for cfDNA batches, colored by sample origin. (H) In the UMAP from (G), three representative oncogenes were marked with warm colors and three representative tumor-suppressor genes in cool colors; these annotations illustrate the model’s separation of oncogenic versus tumor-suppressive contexts. All panels were plotted using training-set data. See Fig. S3 for all cancer types and test-set data validation.

A second model task is tissue-of-origin (TOO) prediction based on the same multi-fragment gene-context. We quantify prediction confidence from the model’s origin logits and observe that batches with high confidence exhibit substantially higher accuracy (Fig. 3B), whereas low-confidence batches tend to produce high-scoring false positives. Applying a simple confidence-index cutoff to select high-confidence batches and multiplying the cancer score by the confidence yields a gated score used for downstream analysis. Gated cancer scores are profoundly different between normal and cancer samples, both within individual cancer types (Fig. 3C) and across all cancer types (Fig. 3D), with significance exceeding p < 1×10−³⁰⁰ (Mann–Whitney U). The per-cancer breakdown shows colorectal cancer (CRC) and hepatocellular carcinoma (HCC) contribute most positive batches, while lung adenocarcinoma (LUAD) contributes few, consistent with its well-documented low ctDNA shedding. The origin-prediction confusion matrix recapitulates this result: CRC is rarely misclassified, whereas many LUAD batches are predicted as healthy (Fig. 3E). Although most genes are not reliably called positive across batches, a minority of genes are stably detected in many batches—for example, ADAMTS9 and ALDOB, consistent with their reported hypermethylation-silencing (Fig. 3F).

Internally, cfAI’s attention mechanisms learn multimodal representations: the summary CLS token progressively aggregates useful signals for downstream tasks. Unsupervised UMAP projection of the CLS hidden states places cfDNA batches from different tissue origins into distinct regions of latent space, producing an integrated atlas that both predicts and clusters sources while retaining cancer-relevant biology (Fig. 3G). Within single-cancer clusters, the model spontaneously segregates groups whose representative genes are enriched for oncogenes versus tumor-suppressor genes, indicating that the model has internally learned different functional roles of these two gene classes.

### Pseudo-labeling strategy for model architecture and training regimen

Building cfAI as a multimodal foundation model with cfDNA as the token necessitated architecture and training strategies tailored to four nested requirements: accurate alignment of multimodal features at the single-cfDNA level, crosstalk among multiple cfDNA tokens, preservation of gene-level context, and learning from labels that are only available at the sample level. Without a reliable ctDNA gold standard, we adopted a pseudo-label, adversarial training paradigm in which all cfDNA fragments derived from cancer samples are initially treated as candidate ctDNA. Given that this label is highly inconsistent with ground truth, and our goal is to recover ≈0.1% true ctDNA from a background of 99.9% non-ctDNA noise, we implemented a mixed multi-label sampling strategy. For each target sample (cancer or normal), we incorporate depth-matched normal background noise, which is sampled from ten randomly selected normal training samples; the model was then trained to identify and recover putative cancer-derived fragments from this mixed batch. This approach prevents entire batches from being uniformly labeled as ctDNA and preserves discriminative learnable contrasts.

Batch construction uses expression-aware, gene-wise sampling to preserve gene-level context. Token batches are generated asynchronously by gene, and we dynamically include one or more similar genes plus padding tokens to assemble a context of appropriate length given the available cfDNA counts. Critically, sampling was scaled by the sample’s total sequencing depth rather than per-gene read counts, allowing extraction of depth differentials between the target and noise background; we embed histone-enrichment proxies derived from these depth differences into the batch CLS token so that gene-internal context and inferred expression are aligned during training (Fig. 4A).The training objective is a weighted combination of three prediction losses: per-cfDNA score, cfDNA gene-batch aggregated score, and origin-type logits. Because the true positive rate (TPR) is non-differentiable, it cannot be used directly as a loss; instead, we monitor TPR as an external metric while optimizing the differentiable surrogate losses.

**Figure 4.**
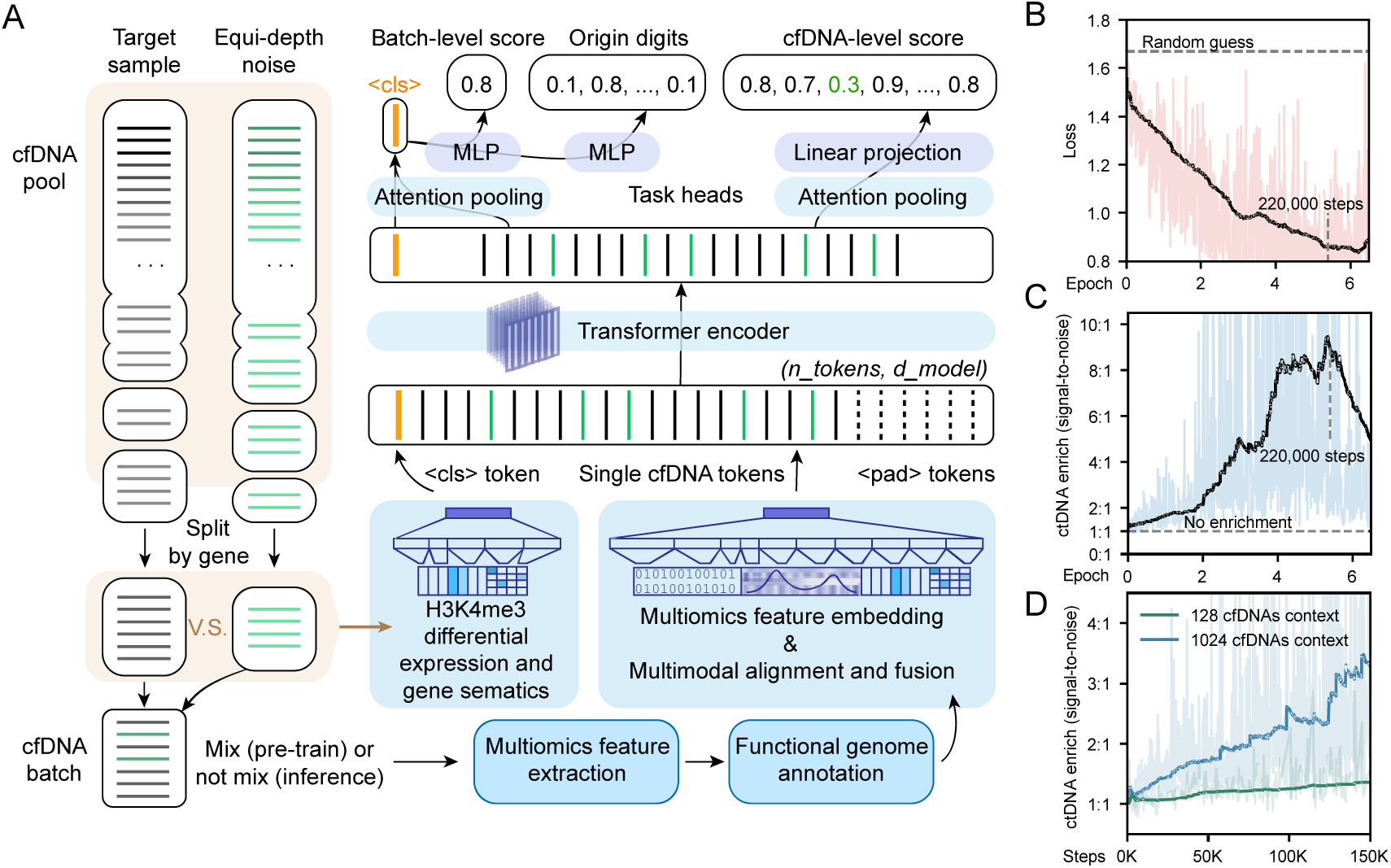
Model architecture and training workflow. (A) Expression-aware gene-wise sampling mixes target and depth-matched normal background to produce pseudo-labeled cfDNA batches; expression metrics (depth ratios and up/down direction) are concatenated with gene semantics into the CLS token and fed together with multimodal cfDNA tokens into a standard Transformer. Attention pooling aggregates token outputs to produce per-cfDNA scores, and a secondary attention pooling produces a CLS embedding for batch-level discrimination. (B–D) Training trajectories for loss and monitoring metrics; the initial random-guess baseline is annotated. Each epoch corresponds to a full pass through the training samples (∼40,000 steps per epoch, with a step defined as one batch). cfDNA enrichment is defined as the ratio of mean cancer-origin cfDNA score to mean noise cfDNA score and quantifies improvement in signal-to-noise.

During training, we observe steady loss reduction accompanied by large improvements in signal-to-noise: the mean score ratio of cancer-origin fragments to noise fragments increases from approximately 1:1 to near 10:1 (Fig. 4B–C). We evaluated a reconstruction pretraining task (reconstructing methylation or other modalities from sequence and orthogonal features) as an easy auxiliary objective to accelerate convergence; although it sped early convergence, it did not materially improve final performance on the principal detection task (data not shown). Hyperparameter and configuration tuning show several practical findings. Lower learning rates allow stable training for more epochs; learning-rate schedulers help escape saddles; and resuming interrupted runs reliably restarts loss descent, with continued improvement observed after ten or more epochs. Context length has the largest impact on discriminative power: 1,024 cfDNA tokens of context substantially outperform 128 tokens for ctDNA enrichment (Fig. 4D). We adopted early stable checkpoints at 220,000 training steps to mitigate overfitting. While the model architecture remained fixed (around 41 million parameters), it can continue to be optimized through repeated training restarts (Fig. S4). In the longest training run, the model was trained for up to 500,000 steps, resulting in a total exposure of approximately 8 billion training tokens (500,000 steps × 16 batch size × 1,024 token context). During inference we retain the same background-sampling and expression-comparison procedures to compute gated scores, but we omit the injected noise tokens when feeding inputs to the Transformer; the model then selects candidate ctDNA fragments from the target tokens.

### Sample-level cancer signal detection

For sample-level detection, we employed the mean of the cfDNA-batch cancer scores as the metric for sample discrimination. Due to the variable number of gene-level cfDNA batches per sample and the limited size of the cohort, we did not develop a complex classifier at the sample level; instead, inference was performed with fixed model parameters, and results were output for analysis. Both the training and test datasets demonstrate clear separation between normal and cancer samples (Fig. 5A), with the area under the ROC curve (AUC) values of 0.845 and 0.887 for the training and test sets, respectively (Fig. 5B). We established a decision cutoff on the training set to attain a specificity of 94.3% (95% CI 84.3–98.8, 50/53) and subsequently applied this cutoff to the held-out test set. On the test set, this cutoff corresponds to a specificity of 93.1% (95% CI 77.2–99.2, 27/29) and an overall sensitivity of 72.6% (95% CI 62.5–81.3, 69/95). Sensitivity of cancer signal detection by cancer type and clinical stage is shown in Fig. 5C and Fig.5D, respectively. The sensitivity of various types of cancer is as follows: colorectal cancer at 73.1%, liver cancer at 84.6%, lung cancer at 70%, and the others at 69.2% (Fig. 5C). Previous studies using the cfDNA methylation biomarker for multiple cancer early detection demonstrated good sensitivity in liver and colorectal cancers, which is consistent with our findings. cfAI achieved a sensitivity of 72% (8/11) for stage I-II colorectal cancer, and 80% (4/5) for stage I liver cancer. Early-stage lung adenocarcinoma is known for typically minimal ctDNA shedding into the peripheral bloodstream, which generally achieves a low sensitivity (< 15%). In this study, our model achieved stable separation from non-cancerous samples (Fig.5D), and the positive detection rate for stage I lung adenocarcinoma was 67% (6/9) in the test set.

**Figure 5.**
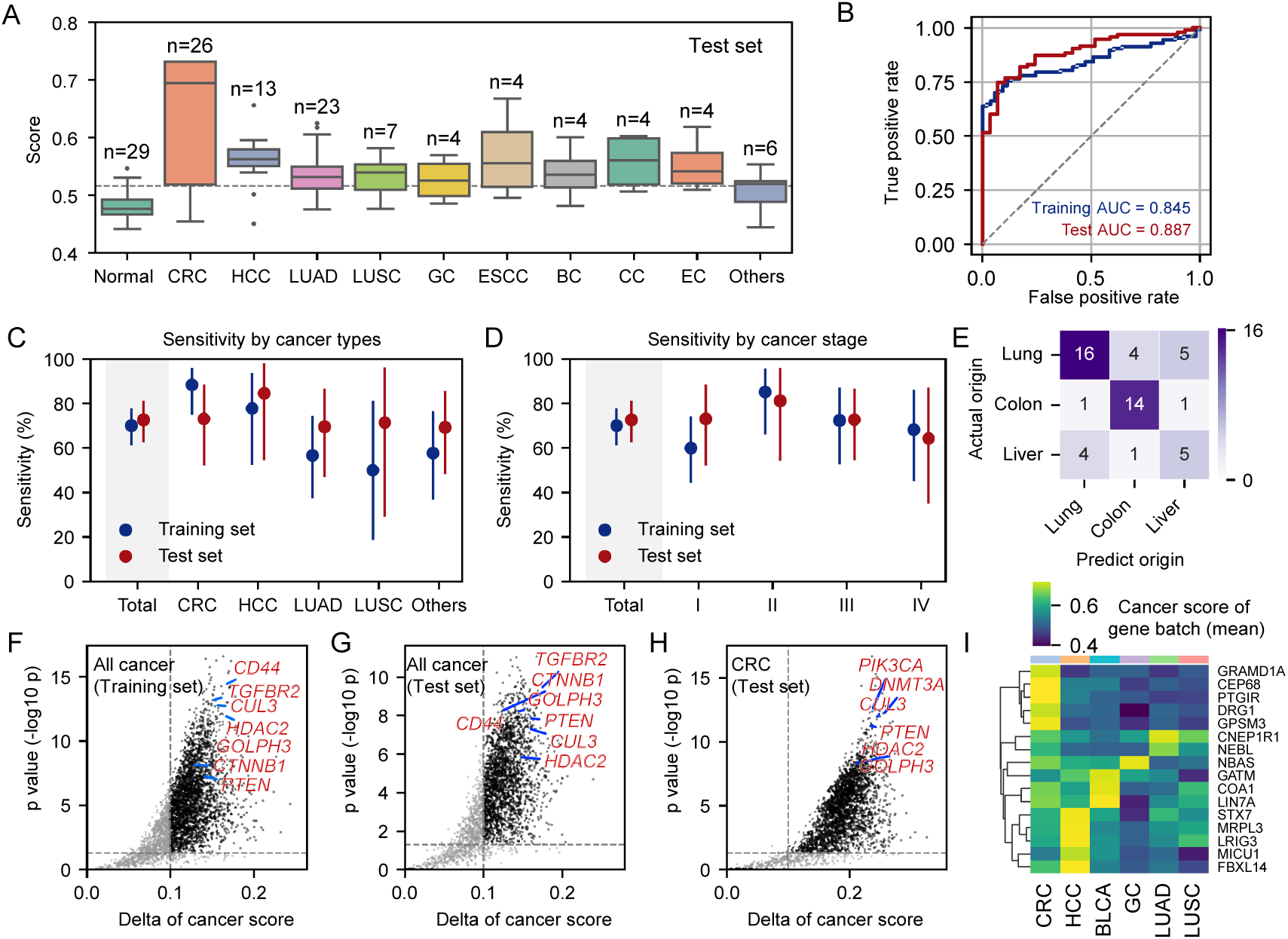
Performance evaluation of cfAI. (A) Box plot of sample diagnostic scores by cancer type in the test set; the dashed line indicates the cutoff derived from the training set (cutoff = 0.5156). (B) ROC curves for training and test sets with AUC annotations. (C–D) Sensitivity by cancer type and by clinical stage in training and test sets. (E) Confusion matrix for sample-origin classification on the test set. (F–H) Gene-level cancer-score differences between cancer and normal samples: (F) all cancer types in the training set, (G) all cancer types in the test set, and (H) CRC in the training set. The x-axis shows score delta, the y-axis shows t-test significance; genes with significant biological relevance are highlighted in red. (I) Relationship of cancer-type–specific genes across tumor types, with color denoting the mean cancer score. See Fig. S5 for results in training set.

TOO prediction at the sample level is impeded by unequal cohort sizes; the model tends to disproportionately assign predictions to CRC and lung cancers, which also exhibit higher per-class accuracy, whereas other cancer types—with fewer samples—are underrepresented in predictions (Fig. 5E). To evaluate each gene’s contributions for diagnosis across cancer types, we performed statistical testing of gene-wise scores and identified multiple biologically meaningful genes that are consistently elevated in both training and test sets (e.g., HDAC2, TGFBR2) (Fig. 5F–G). CRC shows enrichment of genes in pathways including PI3K and DNMT3A (Fig. 5H). We also identify cancer-type–specific candidate markers such as DRG1 for CRC, which may serve as useful leads for future development of cancer-specific markers.

## Discussion

This study advances cfDNA analysis by both increasing resolution and preserving the full complexity of multi-omic interactions. Prior work^29,30^ identified pairwise co-occurrences on individual fragments—e.g., methylation paired with fragmentomics or SNVs paired with methylation—but analysis remained dominated by selective screening and mixture-level statistics; such approaches achieve an improvement analogous to the left middle transition in Fig. 2B. In reality, cfDNA carries far richer and more heterogeneous co-signals that vary by gene and genomic position, and simple filtering or categorical selection both reduces the number of retained fragments and severs interactions among classes, making rare true signals progressively harder to recover. By contrast, our framework maintains every cfDNA feature while modelling high-order, multi-omic cross-interactions—effectively enabling full-factorial inference at extremely fine granularities.

Delivering that capability requires a purpose-built multimodal foundational model and a domain-specific semantic system for omics. Off-the-shelf large language and vision models do not natively encode the scalar, periodic, and contextual patterns characteristic of omic modalities, and single-cell or gene-level models lack a tokenization schema that treats individual cfDNA molecules as first-class inputs. We therefore developed, from first principles, an end-to-end multimodal cfDNA foundation model whose learnable embeddings and attention mechanisms jointly encode molecular measurements together with genomic context; this architecture not only enriches ctDNA but also internalizes biologically meaningful distinctions such as oncogene versus tumor-suppressor behavior.

The study also addresses a key practical limitation of prior screening efforts: the tradeoff between model complexity and available sample counts. Conventional approaches face a parameter-vs-sample “curse” that constrains model scale^37–39^. By treating cfDNA and gene-level cfDNA batches as abundant training signals, we expand effective training volume from industry-typical thousands to the billion-scale token regime, enabling larger models that can learn subtle cross-modal structure without overfitting. To capture more orthogonal signals in the proof-of-concept, we used histone-modification enrichment to focus sequencing on promoter-proximal regions where cancer signals concentrate; this reduces training burden and concentrates discriminative features while adding one informative modality.

That experimental choice, however, carries practical caveats. Antibody pull-down is inherently variable, and performing ChIP-like (Chromatin ImmunoPrecipitation) enrichment directly in plasma is sensitive to preanalytical variables such as storage conditions; these factors complicate engineering and commercialization. Crucially, the analytical strategy is transferable: the same embedding and fusion paradigm can be applied to whole-genome sequencing or targeted capture panels and to non-ChIP enrichment schemes now favored in clinical pipelines (e.g., enzymatic methylation conversion), allowing fragmentomics and methylation to be jointly modelled without relying on plasma ChIP.

There are several limitations of this study. Due to the limited sample size, the current model training is not sufficient. Therefore, it is speculated that there is still room for improvement in its performance. Additionally, it is currently challenging to refine the performance for each stage. The accuracy of TOO prediction was slightly lower in this study than that reported in other published large-sample studies, which can be attributed to the limited sample size used for model training. Even so, during the preliminary model analysis, it was observed that different types of cancer reads could be well distinguished and clustered, reflecting the model’s prediction potential in TOO. For cfAI to be used in the early detection of multiple cancers in the future, TOO is a key indicator for the diagnosis, which still needs to be trained and validated with a larger number of samples.

In conclusion, this study demonstrates a single-molecule, multimodal fusion of cfDNA and delivers an unprecedented degree of information integration. By preserving every fragment’s feature set and modelling their complex cross-interactions, the approach yields new mechanistic insight and a practical blueprint for next-generation, noninvasive liquid biopsies and early cancer screening.

## Methods

### Study design and cohort

This observational case–control study prospectively recruited a total of 333 participants (cancer patients across 10 cancer types and non-cancer controls), between August 2022 and November 2024, from 11 medical sites. All participants provided written informed consent. For every participant, demographics, general clinical information, medical history, and cancer diagnosis data were collected. The clinical stages of cancers were defined according to the American Joint Committee on Cancer 8th edition. After plasma and sequencing quality control, 304 samples qualified for analysis and were randomly split, after matching for age, sex and cancer type, into training (n = 180) and testing (n = 124) sets (Fig. S2). Detailed clinical characteristics and inclusion/exclusion criteria are provided in Table S1 and Table S2.

### Blood collection and plasma preparation

Peripheral blood (8 mL) was collected into K3-EDTA tubes and kept on ice. To limit loss of histone modifications, samples were supplemented immediately with 1× protease inhibitor cocktail (cOmplete™, Roche) and 10 mM EDTA. Plasma was isolated by two centrifugation steps (1,500 × g, 10 min; then 3,000 × g, 10 min; both at 4 °C) and aliquoted for −80 °C storage until processing.

### Antibody-bead conjugation and cf-nucleosome capture

Recombinant anti-Histone H3 (tri-methyl K4) antibody (Abcam, ab213224) was covalently coupled to epoxy-activated Dynabeads (Invitrogen, #14301) using ammonium-sulfate–mediated coupling (≈1 M final [NH₄]₂SO₄) at a 100:1 bead:antibody mass ratio. After overnight coupling with gentle rotation, beads were washed and stored in PBS (0.01% sodium azide) at 4 °C. For chromatin immunoprecipitation, 1 mL thawed plasma was supplemented with inhibitors, incubated with antibody-conjugated beads (2 μg antibody per sample) at 4 °C for 16 h with end-over-end rotation, then washed sequentially in low-stringency Triton buffer and Tris buffer to reduce nonspecific binding^11^. Chromatin was eluted at 55 °C in an SDS/proteinase K buffer, purified with 1.4× SPRI (AMPure XP), and eluted in 21 μL IDTE buffer.

### Library construction and multi-omic capture

Eluted cfDNA underwent nick repair by Taq Ligase (NEB, M0208) and dual-strand library construction, followed by enzymatic methylation conversion using a NEBNext Enzymatic Methyl-seq (EM-seq) workflow (NEB, E7120L) with unique dual indices (E7140L). This protocol preserves fragment-end integrity in highly nicked cfDNA and protects 5mC near fragment termini, guided by our prior observation that nick-repair enzymes such as Taq ligase can largely compensate for 3′ 5mC loss introduced by end-repair polymerases during double-strand EM-seq of cfDNA, reflecting the extensively nicked state of circulating DNA (see Fig. S1 and Supplementary Methods). A 0.02 ng dual-spike-in control (300 bp mixture of CpG-methylated pUC19 and unmethylated λ DNA) was added prior to end-repair. End-prep, adaptor ligation, and SPRI cleanups followed manufacturer recommendations. Conversion/deamination steps used APOBEC/TET-based enzymatic procedures and libraries were PCR-amplified (Q5U polymerase, 15 cycles). Final libraries were purified, assessed on a Bioanalyzer, quantified by qPCR, pooled equimolarly, and sequenced (Illumina NovaSeq 6000, 150 bp paired-end) to an average yield of ∼6 Gb per sample. This protocol preserves fragment end integrity in highly nicked cfDNA and protects 5mC near fragment termini (see Fig. S1 and Supplementary Methods).

### Primary processing and per-read annotation

Raw FASTQ files were processed in a reproducible Nextflow pipeline including steps below. Adapters and low-quality bases were trimmed with Trim Galore! (v0.6.6)^40^. Reads were aligned to GRCh38 with Bismark (v0.22.3)^41^ using seed settings (–N 1 –L 20) and only uniquely mapped reads were retained. PCR duplicates were removed using Bismark’s deduplication. Per-base CpG methylation calls were obtained by the Bismark methylation extractor (–paired-end, –no_overlap). BAM files were used to generate per-read tables by intersecting read coordinates with transcription start site windows (TSS ± 8192 bp), ChromHMM segmentation, A/B compartment annotations, and other regulatory tracks.

### Feature extraction and normalization

For each cfDNA fragment we extracted: nucleotide sequence (A,T,C,G), CpG methylation calls (M/U), mismatch events, per-read methylation ratio (methylated / [methylated + unmethylated]), fragment length, and end-motif sequences. Using genomic coordinates, each fragment was annotated with nearest gene identity, distance to TSS, ChromHMM state, A/B compartment, Hi-C derived features, and other context annotations. Feature-specific normalizations were applied (e.g., TSS distance scaled to 8,192; ChromHMM UMAP embeddings scaled to [–1,+1]; Hi-C values divided by 20) prior to embedding. Custom preprocessing scripts appended sample metadata and computed per-read statistics for downstream modeling.

### Modality-specific embeddings and tokenization

Because cfDNA multi-omics spans categorical, numeric and structured inputs, we implemented modality-aware mappers to convert raw features into semantic vectors:

- Sequence & methylation: One-hot encoding for bases (A/T/C/G), CpG states (M/U) and mismatch markers (E) was input to a 4-layer 1D CNN to produce sequence embeddings.
- Nucleosome/positional features: Fragment length and TSS-relative distance were embedded with learnable Fourier feature encodings initialized to biologically informed wavelengths (DNA helix turns and nucleosome repeat lengths).formulated as:

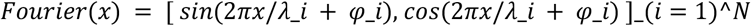 Where λ_i are wavelengths spanning biochemical periodicities.
- End motifs: End-motifs were decoded through an F-profile deconvolution into a 6-dimensional DNase-contribution vector.
- Chromatin states and priors: ChromHMM categorical states used a trainable lookup table; emission-matrix priors^35^ were UMAP-reduced and included as fixed vectors.
- Gene semantics: Gene identity and function were embedded using pretrained scGPT gene embeddings^34^.
- Other quantitative features: Methylation ratios, Hi-C vectors and topological values (IS, DI, FI, AB)^42^ were projected via learnable MLPs.

All per-feature embeddings were concatenated and passed through a projector MLP to yield a 512-dimensional latent vector per cfDNA fragment. Each fragment token therefore preserves the full multi-omic profile and genomic context.

### Transformer architecture, batch construction and training strategy

Tokens were grouped into cfDNA-batches by gene; each batch was prefixed with a <cls> token that encodes batch-level metrics (depth and relative enrichment versus background, gene sematic). During pretraining, target tokens were mixed with noise tokens sampled from a matched-depth pool of healthy cfDNA to simulate realistic background. The classic Transformer encoder (L=6 layers, H=8 heads) processes batches to learn both intra-token representations and inter-token interactions via attention. The model was trained end-to-end with a multi-task loss including token-level ctDNA prediction, batch-level prediction of cancer diagnosis and cancer-type classification digits of this target sample. Training used mixed-precision (AMP), gradient scaling^43,44^, and a cosine annealing learning-rate schedule with warm restarts^45^.

### Inference and downstream classifiers

At inference, samples were run without noise mixing. Sample scores were computed by aggregating cfDNA-batch scores (mean of cfDNA-batch cancer probabilities multiply by cancer-type classification confidence) to produce a final diagnostic score thresholded using cutoffs derived from the training set. Software, reproducibility and data availability

Primary pipelines and custom preprocessing scripts were implemented in Nextflow and Python; key tools include Trim Galore!, Bismark, and custom embedding modules in PyTorch. Versioned code, parameter settings and annotation data are provided in the Supplementary Materials and will be made available upon publication.

## Contributions

Conception and design: Liyang Song, Bo Wang, Xiaohui Wu, Xueguang Sun, Zhidong Gao and Yingjiang Ye

Acquisition of data (participant enrollment, blood samples and clinical information collection, etc.): Bo Wang, Hefei Li, Nan Lin, Xiaowen He, Wenxin Liu, Li Liu, Jian Cui, Xuesong Li, Ying Mei, Qiuting You, Haodong Zhu

Experimental performance : Liyang Song

Development of analysis methodology: Liyang Song

Figure preparation: Liyang Song

Analysis and interpretation of data: Liyang Song, Ying Xin, Guoqiang Zhao

Technical support: Jing Liu, Guoqiang Zhao, Xueguang Sun

Writing, review and/or revision of the manuscript: Liyang Song, Ying Xin, Bo Wang, Guo Chen, Jing Liu, Baoliang Zhu, Xiaohui Wu, Zhidong Gao, Yingjiang Ye

Study supervision: Xueguang Sun, Xiaohui Wu, Zhidong Gao and Yingjiang Ye

## Acknowledgements

We gratefully thank the patients, volunteers and their families for participating in this study. Funding for this study was supported by Shanghai Xiaohe Medical Laboratory Co., Ltd.

## Data and Code Availability

The data supporting the findings of this study are available on request from the corresponding author. All requests for raw and analytical data will be promptly reviewed by trial organizer to determine if there are any intellectual property or confidentiality restrictions. Any data that can be shared will be released via a data use agreement. The Analysis code is available at repositories that will be released recently.

