## Supplementary information for "Multimodal AI for Single cfDNA Profiling and Cancer Screening"

### Supplementary Figures

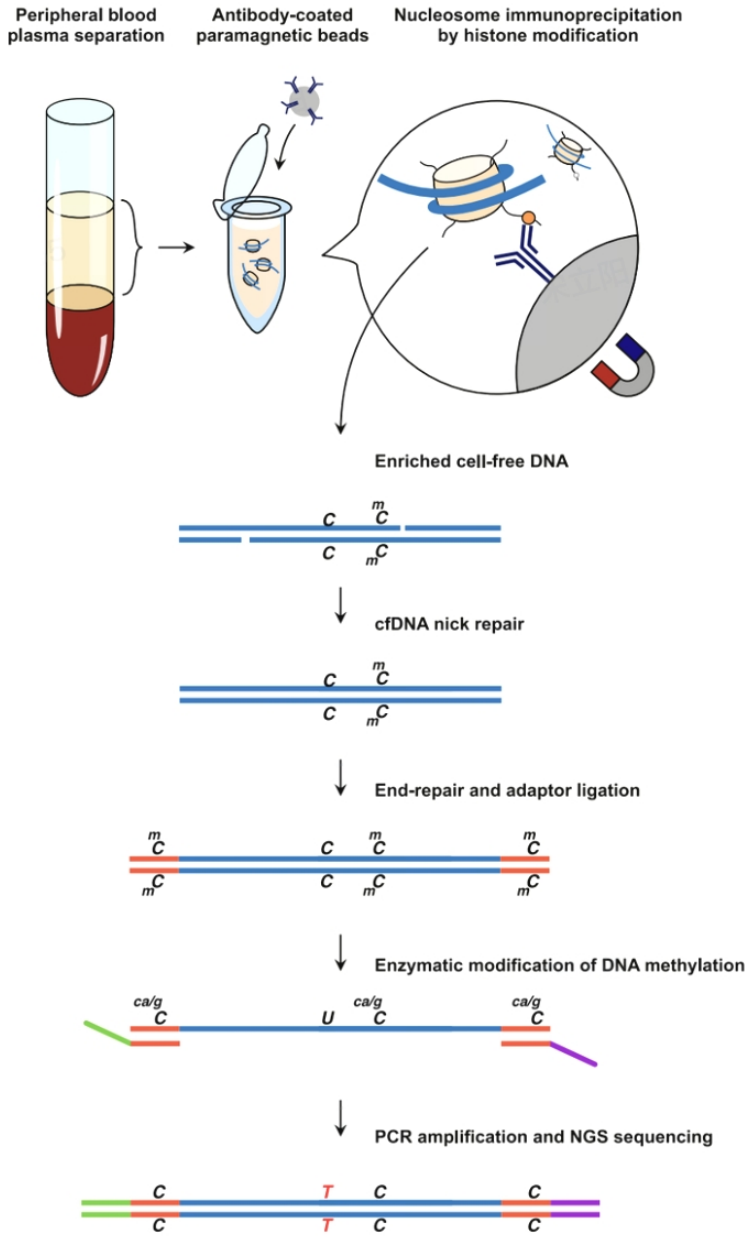


**Supplementary Figure 1. Library preparation workflow.** After capture of cf-nucleosomes, cfDNA is eluted, purified, and repaired, followed by double-strand library construction. Methylation conversion is performed enzymatically using TET2 and DNMT1 to generate sequenceable bases. The library is PCR-amplified and sequenced on a next-generation sequencing instrument.

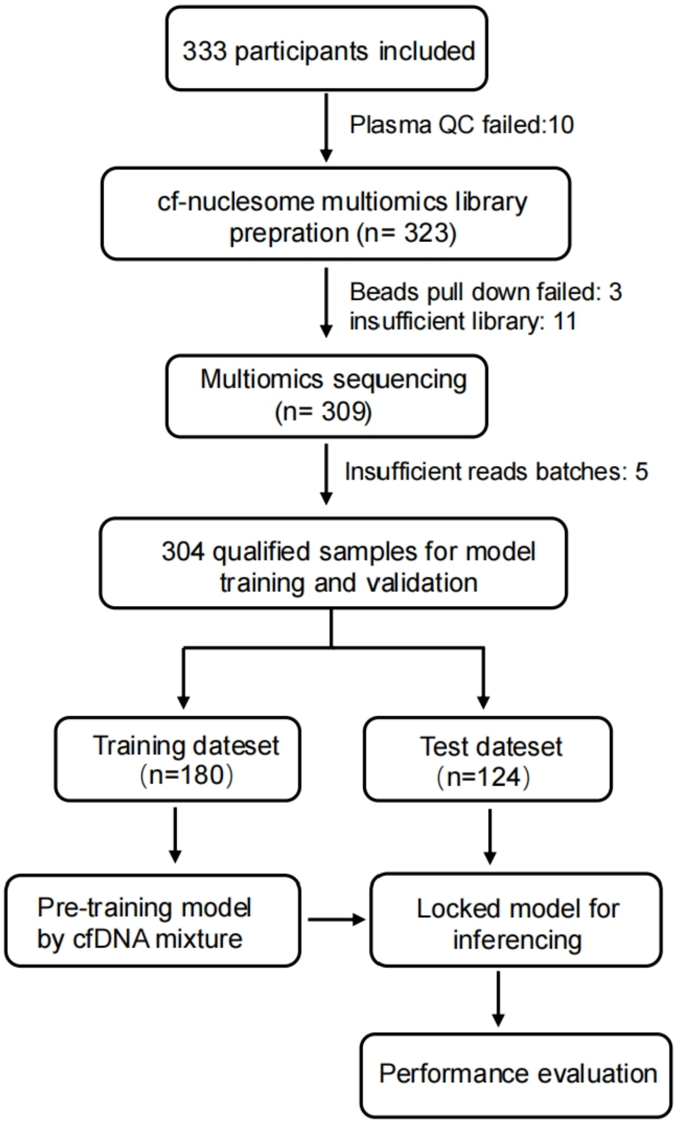


**Supplementary Figure 2. Flow diagram of participant disposition.** The flow diagram illustrates prospective recruitment, sample quality control, and cohort allocation for model training and locked test-set evaluation.

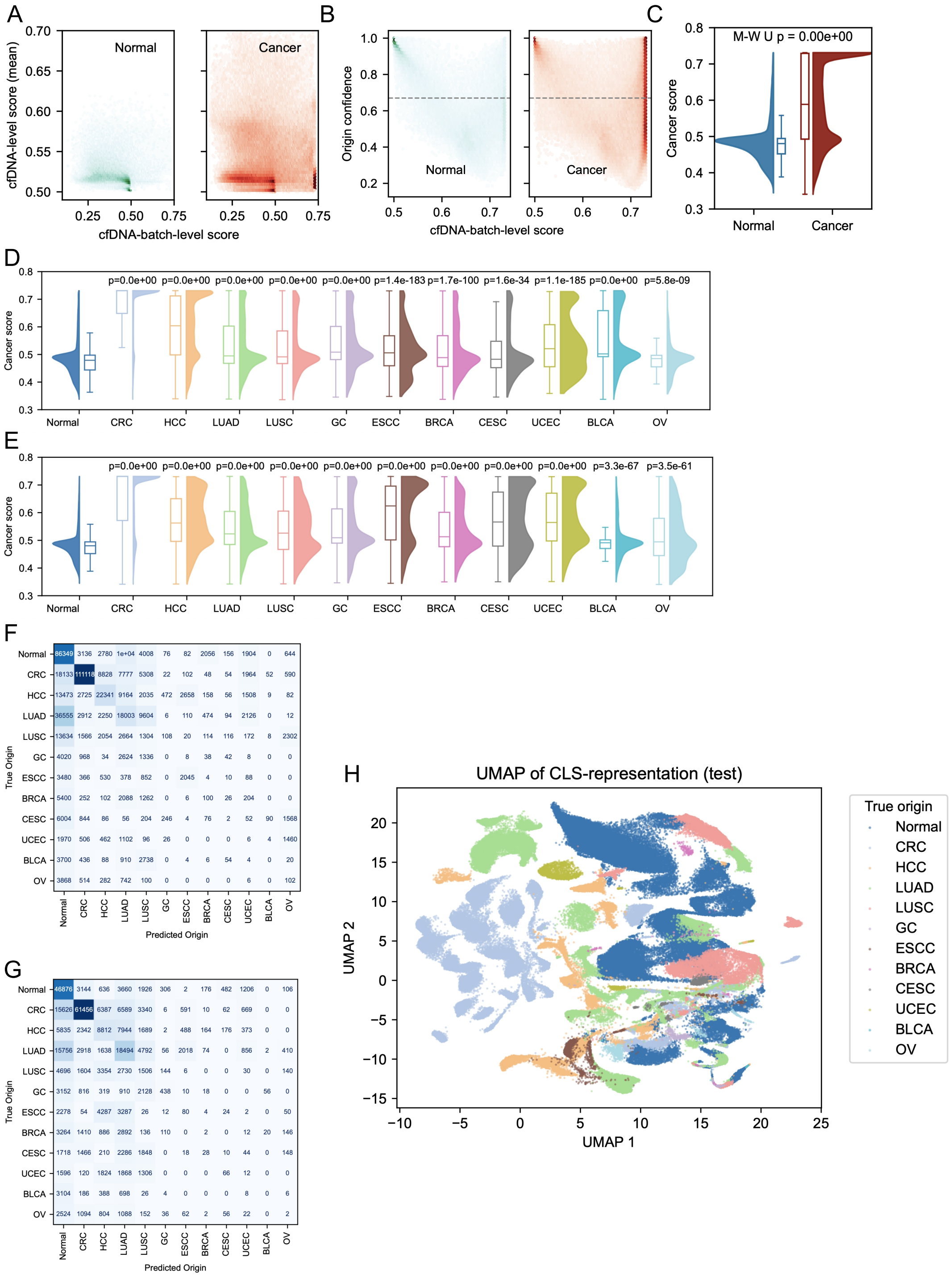


**Supplementary Figure 3. Test-set validation of contextual denoising and discrimination.** Panels correspond to Figure 3, with test-set results shown, unless (D, F), training-set results across all cancer types.

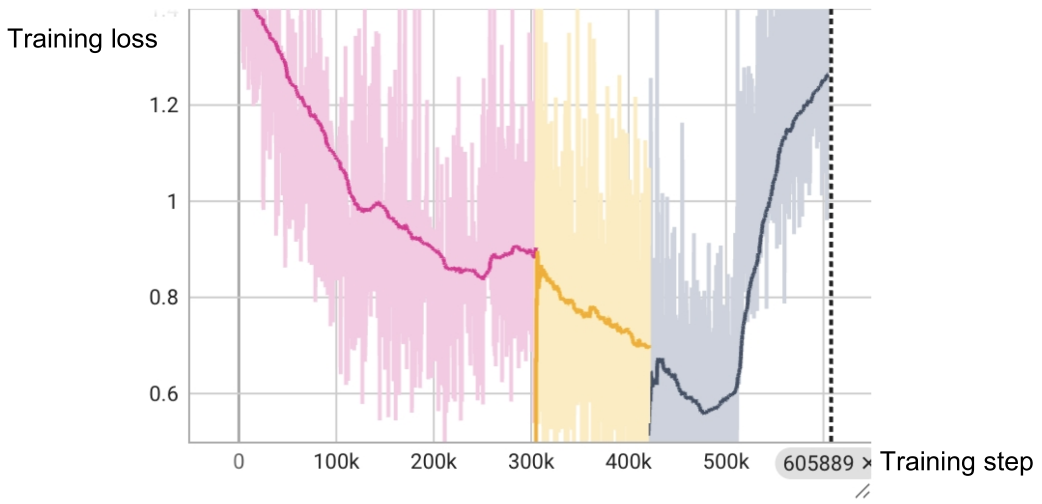


**Supplementary Figure 4. Training loss trajectory under continued optimization.** After fixing the model architecture at step of 220k, training was resumed multiple times from stable checkpoints (different color). With increasing total training steps, the loss steadily decreased, reflecting improved convergence. Transient loss spikes were mitigated by checkpoint rollback and restart.

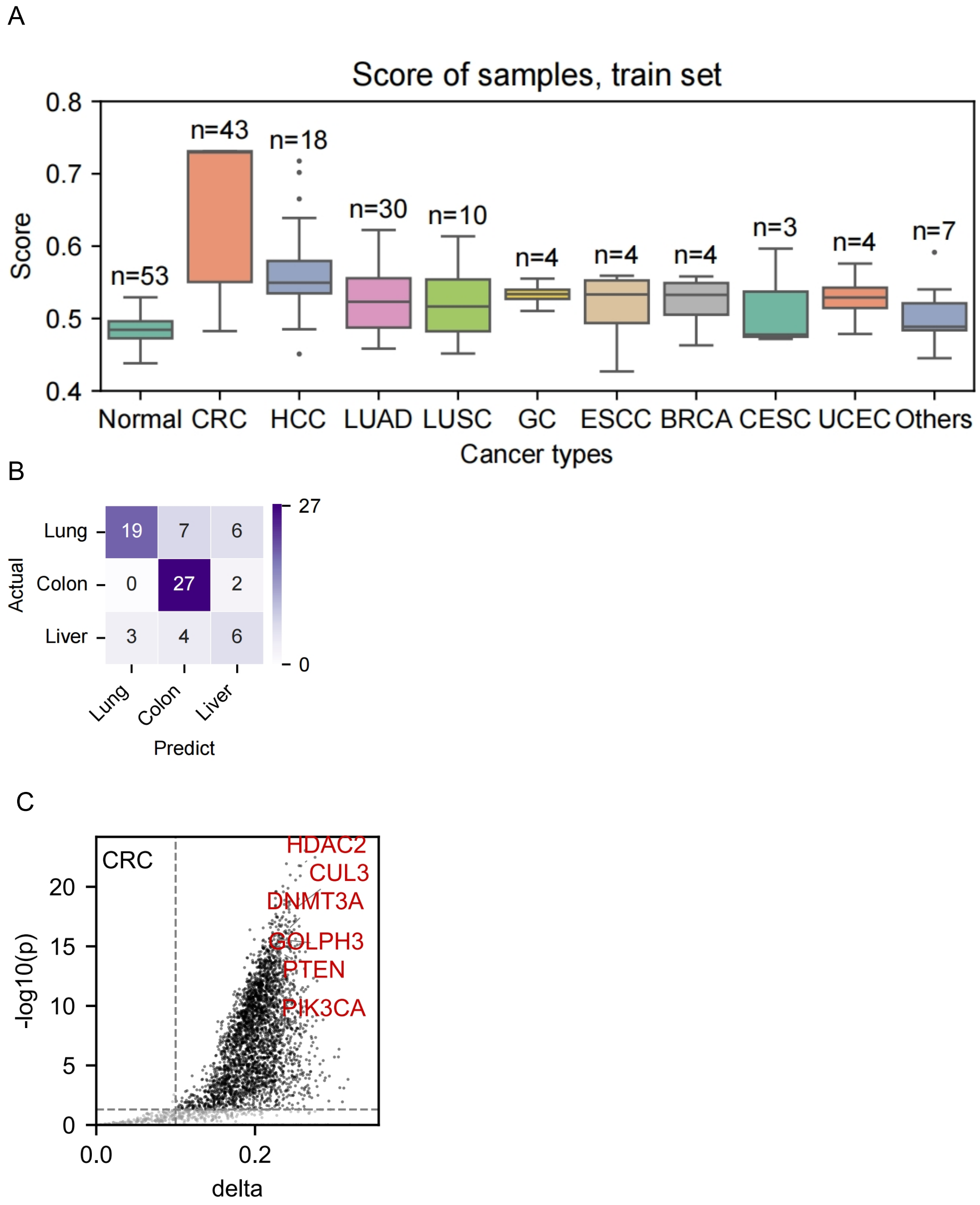


**Supplementary Figure 5. Sample-level diagnostic performance in training set.** Panels correspond to Figure 5 A, E and H, with training-set results shown.

### Supplementary Tables

| Clinical characteristics of study participants | | | | |
|---|---|---|---|---|
| | Training Cancer （n=127） | Training Non-cancer （n=53） | Test Cancer （n=95） | Test Non-cancer （n=29） |
| **Age，median（IQR)** | 63（57-68） | 59（52-65） | 60（53-69） | 59（49-66） |
| **Gender，n (%)** | | | | |
| Male | 57（44.9） | 22 (41.5) | 49（51.6） | 12（41.4） |
| Female | 70（55.1） | 31(58.5) | 46（48.4） | 17（58.6） |
| **TNM stage，n (%)** | | | | |
| Tis | 1 (0.8) | | 2 (2.1) | |
| I | 45 (35.4) | | 26 (27.4) | |
| II | 27 (21.3) | | 16 (16.8) | |
| III | 29 (22.8) | | 33 (34.7) | |
| IV | 25 (19.7) | | 18 (18.9) | |
| **Cancer type, n (%)** | | | | |
| Colorectal cancer | 43 (33.9) | | 26 (27.4) | |
| Lung cancer | 40 (31.5) | | 30 (31.6) | |
| Liver cancer | 18 (14.2) | | 13 (13.7) | |
| Breast cancer | 4 (3.1) | | 4 (4.2) | |
| Endometrial cancer | 4 (3.1) | | 4 (4.2) | |
| Esophageal cancer | 4 (3.1) | | 4 (4.2) | |
| Gastric cancer | 4 (3.1) | | 4 (4.2) | |
| Ovarian cancer | 4 (3.1) | | 3 (3.2) | |
| Cervical cancer | 3 (2.4) | | 4 (4.2) | |
| Urothelial cancer | 3 (2.4) | | 3 (3.2) | |

**Supplementary Table 1. Clinical characteristics of study participants.**
Summary statistics of participants and clinical information, including age, sex, disease status, and group distribution.

| | Inclusion Criteria | Exclusion Criteria |
|---|---|---|
| All | 1. Age：30~80<br>2. Able to provide a written informed consent | 1. Pregnancy or lactation.<br>2. A history of severe acute infectious diseases within the past 14 days before blood draw.<br>3. Use of the following medications within the past 14 days: intravenous/oral glucocorticoids; intravenous/oral azacitidine, decitabine, procainamide, hydralazine, or arsenic trioxide.<br>4. A history of organ transplantation, allogeneic bone marrow transplantation, or stem cell transplantation, or receipt of blood transfusion within 30 days before blood draw.<br>5. Other circumstances deemed unsuitable for participation in this study by the investigator. |
| Cancer | 1. Confirmed cancer diagnosis based upon assessment of a pathological specimen or a high suspicion for a cancer diagnosis by clinical and/or radiological assessment, with planned biopsy or surgical resection to establish a definitive diagnosis within 4 weeks after blood draw.<br>2. Has not received any systemic or local anti-tumor treatment, including but not limited to surgical resection, radiotherapy, chemotherapy, hormone therapy, targeted therapy, immunotherapy, interventional therapy, etc. | |
| Non-cancer | 1. Self-reported absence of any tumor history, and clinically judged by a physician to be free of any signs of tumors.<br>2. Diagnosed clinically and/or pathologically with a benign disease (e.g.benign pulmonary nodules) | |

**Supplementary Table 2. Clinical inclusion/exclusion criteria.**

Summary of inclusion/exclusion criteria for cancer and non-cancer participants.

### Supplementary Methods

**Reagents and Equipment**

Key reagents included anti-H3K4me3 antibody (Abcam, ab213224), epoxy-activated Dynabeads M270 (Invitrogen, 14301), Amicon Ultra-0.5 mL 10 kDa filter units (Millipore, UFC501024), cOmplete™ protease inhibitor tablets (Roche, 4693132001), and NEBNext Enzymatic Methyl-seq library prep kits (E7120L/E7140L). All centrifugation steps utilized a 4 °C pre-cooled benchtop centrifuge. Bead couplings were performed in LoBind tubes with constant end-over-end mixing.

**Antibody Buffer Exchange**

To remove glycerol and azide, 500 µL of antibody solution was buffer-exchanged into PBS (pH 7.4, 0.02% sodium azide) using an Amicon Ultra-0.5 filter (10 kDa cutoff). Samples were spun at 14,000 × g for 5 min, diluted to volume, and repeated twice. The final concentrate was adjusted to original volume with PBS.

**Dynabeads Preparation**

On receipt, dry beads were equilibrated to room temperature, resuspended at 30 mg/mL in anhydrous DMF, and stored at 4 °C. Prior to coupling, beads were washed twice with phosphate buffer (0.1 M sodium phosphate, pH 7.4) to remove DMF.

**Coupling Reaction**

For each sample, 5 mg beads were pelleted and washed in buffer A (0.1 M phosphate, pH 7.4). Beads were resuspended in 100 µL buffer A, then 100 µL antibody (0.5 mg/mL) and 100 µL 3 M ammonium sulfate in buffer A were added (final $[NH_4]_2SO_4 \approx 1$ M). The 300 µL reaction was incubated at room temperature (≈22 °C) with gentle rotation for 16–24 h. Post-coupling, beads underwent four PBS washes and were finally resuspended in PBS containing 0.01% sodium azide.

**cf-nucleosome capture Enrichment**

1 mL plasma was thawed on ice, supplemented with 40 µL 25× protease inhibitor storage solution and 20 µL 0.5 M EDTA. Beads (200 µg, ~6.6 µL) were added, and mixtures were rotated at 4 °C for 16 h. After incubation, beads were magnetically separated; unbound plasma was aliquoted and frozen. Beads were washed four times with Blood Wash Buffer (50 mM Tris-HCl pH 7.5, 150 mM NaCl, 1% Triton X-100, 0.1% deoxycholate, 2 mM EDTA) containing 1× protease inhibitors, followed by two washes in 10 mM Tris-HCl pH 7.4.

**Elution and Cleanup**

Beads were resuspended in 50 µL Elution Buffer (10 mM Tris-HCl pH 8.0, 300 mM NaCl, 0.6% SDS, 5 mM EDTA) supplemented with 2.5 µg proteinase K and incubated at 55 °C for 1 h. The supernatant was recovered and purified using 1.4× AMPure XP beads; libraries were eluted in 21 µL IDTE.

**Library Construction and Quality Control**

Libraries were prepared using the NEBNext Enzymatic Methyl-seq Kit with unique dual indices (E7140L), following standard cycling conditions (4 min initial end repair, 4–6 min adaptor ligation, 4 rounds of indexing PCR). Library size distribution was assessed on a Bioanalyzer; concentration was quantified by qPCR. Prepared libraries were stored at –20 °C until sequencing.

#### Detailed Library Preparation and Sequencing Protocol

**Spike-in Preparation ("Dual-spike-300 bp")**

Mix 2.4 µL CpG-methylated pUC19 (0.1 ng/µL) and 2.4 µL unmethylated λ DNA (2 ng/µL) in 1× TE to 130 µL.

Fragment to ~300 bp on a Covaris (microTUBE AFA Fiber Pre-Slit Snap-Cap).

Purify with 1.0× AMPure XP; elute in 240 µL EM-seq Elution Buffer.

Measure concentration (~0.02 ng/µL), dilute to 0.002 ng/µL, aliquot, store at –20 °C.

**Adapter Sequences**

5′–A5mCA5mCT5mCTTT5mC5mC5mCTA5mCA5mCGA5mCG5mCT5mCTT5mC5mCGAT5mC* T–3′

5′–[Phos]GAT5mCGGAAGAG5mCA5mCA5mC GT5mCTGAA5mCT5mC5mCAGT5mCA–3′

**Ligation & End-Prep (Off-Bead)**

**Ligation Pre-mix:** to 21 µL eluate add Taq ligase buffer (2.5 µL), dual spike (0.5 µL, 1:2.5 pre-dilution), Taq DNA ligase (1 µL); incubate 45 °C for 15 min, hold at 4 °C.

**End-Repair:** add NEBNext Ultra II End Prep Reaction Buffer (3.5 µL) and Enzyme Mix (1.5 µL); incubate 20 °C 30 min → 65 °C 30 min → 4 °C.

**Adaptor Ligation:** pre-mix 15 µL Ligation Master Mix + 0.5 µL Ligation Enhancer on ice; add 15.5 µL to sample, then 1.25 µL EM-seq adaptor (0.4 µM); incubate 20 °C 15 min → 4 °C.

**Cleanup:** 1.4× AMPure XP (65 µL); elute in 17.5 µL EM-seq Elution Buffer; store at –20 °C.

**Conversion, Deamination & PCR**

**TET Oxidation & Elution:** add 13.5 µL TET mix; incubate per kit.

**NaOH Denaturation:** add 1.5 µL 0.2 N NaOH; 50 °C 10 min; quench.

**APOBEC Deamination:** add 25 µL APOBEC mix; 37 °C 3 h.

**PCR Setup:** directly add 4 µL EM-seq Index Primer + 44 µL Q5U Master Mix; total 90 µL.

**Cleanup:** 0.9× SPRI; elute in 15 µL TE.

**PCR Cycling:** 98 °C 30 s; 15× (98 °C 10 s, 62 °C 30 s, 65 °C 60 s); 65 °C 5 min; 4 °C hold.

**QC, Pooling & Sequencing**

Assess library size on Bioanalyzer; quantify by qPCR.

For 96 samples, stratify by yield into eight concentration bins (A–H).

Pool equimolarly within bins; perform additional 1.0× SPRI cleanup.

Sequence on NovaSeq 6000 (150 bp PE) aiming for 6 Gb per library; low-yield bins (A–C) targeted at 12–25 Gb to ensure ≥100K spike-in reads.

#### Supplementary Methods: Sequencing Data Analysis

**Workflow Management**

Analysis was orchestrated with Nextflow (v20.10.0), ensuring reproducibility and parallel execution on an HPC cluster. The pipeline definition is available at <repository URL>.

**Read Preprocessing**

**Adapter and Quality Trimming:** Trim Galore! v0.6.6 with default parameters.

**Quality Control:** Post-trim read quality assessed by FastQC v0.11.9.

**Alignment and Methylation Calling**

**Alignment:** Bismark v0.22.3; Bowtie2 backend; options --score_min L,0,-0.2 --N 1 --L 20 to balance sensitivity and specificity.

**Deduplication:** bismark --deduplicate to remove PCR duplicates.

**Methylation Extraction:** bismark_methylation_extractor --paired-end --no_overlap --gzip, producing per-CpG coverage and methylation proportion files.

**Per-Read Annotation and Normalization**

Raw per-read feature tables (columns: M, U, dist_TSS, chromHMM_UMAPemb_1–4, is_value, di_value, fi_value, ab_value, hic_matx/y/z, hic_fatx/y/z) were processed as follows:

a. **Filtering**

Remove any row containing missing entries ("", "NaN", or ".").

Retain reads with dist_TSS within ±8,192 bp.

b. **Metadata Addition**

Append sample_id, origin, and diagnosis fields from the sample manifest.

c. **Feature Computations**

**Methylation Ratio (meth_ratio):** number of "1"s in M divided by sum of "1"s in M and U (set to 0 if denominator = 0).

**Distance to TSS (dist_TSS):** divided by 8,192 to normalize to [–1, +1].

**ChromHMM Embeddings:** each of the four UMAP coordinates multiplied by 2 and subtracted by 1 (rescaling to [–1, +1]).

**Binary Features:**

is_value shifted by –1 (so that original values [1,2] map to [0,1]).

di_value divided by 36.

fi_value shifted by –1.

ab_value left unchanged.

**Hi-C Features:** all six Hi-C distance and contact frequency columns divided by 20.

**Implementation Details**

The entire filtering and normalization workflow was implemented in a single awk one-liner wrapped in SLURM job scripts, ensuring line-by-line stream processing for minimal memory footprint.

Processed tables (.txt) were output to the model-input directory, with automatic logging of input and output line counts for QC.

**Downstream Use**

The finalized per-read feature tables formed the training and testing inputs for our predictive models, as described in the Results section.
